# Modelling Predictive Coding in the Primary Visual Cortex (V1): Layer 2/3 Circuits for Prediction Error Computation through Compartmentalized Spiking Neurons

**DOI:** 10.1101/2025.11.01.686040

**Authors:** Elnaz Nemati, Catherine E. Davey, Hamish Meffin, Anthony N. Burkitt

## Abstract

Cortical Layer 2/3 has been consistently implicated as the locus of prediction-error signalling in hierarchical models of cortical sensory processing. However, the circuit mechanisms that generate biologically plausible prediction-error (PE) signals remain elusive. A spiking network model is presented here in which two-compartment excitatory pyramidal neurons interact with three inhibitory subtypes: parvalbumin-expressing (PV), somatostatin-expressing (SOM), and vasoactive-intestinal-peptide-expressing (VIP) interneurons, to compute sign-specific prediction errors (positive and negative PEs). Feedforward input targets the soma, whereas top-down feedback reaches the distal apical dendrite, enabling a local somato-dendritic comparison. A PE emerges whenever the balance between excitation and inhibition is selectively disrupted within one compartment, recruiting either positive-error (PE^+^) or negative-error (PE^−^) subpopulations of pyramidal neurons. Unlike prior learning-dependent, rate-based accounts, this fixed-weight spiking circuit shows that bidirectional PE signals (PE^+^ and PE^−^) can arise online from compartment-specific balance without any synaptic weight updates. The model reproduces key experimental observations, including sparse mismatch responses, compartment-specific inhibition, and VIP-mediated disinhibition. Across four canonical sensory–prediction configurations, the circuit maintains a tight balance during matched input and generates bidirectional PE signals only under mismatch. By routing sensory drive from Layer 4 into Layer 2/3 and allowing the resulting PE activity to project toward deeper feedback generators, the model situates Layer 2/3 as a dedicated, feature-specific mismatch detector within a hierarchical inference network. These results provide a mechanistic bridge from dendritic computation to laminar predictive coding, demonstrating how realistic spiking dynamics can implement fast, sign-specific PE signaling without learning.

**Author summary:** In this study, we present a biologically grounded spiking model of layer 2/3 in primary visual cortex within the predictive coding framework. Our goal is to explain how superficial cortical circuits compute fast, sign-specific prediction errors when sensory input does not match top-down expectations. The model uses two-compartment pyramidal neurons whose somata receive feedforward drive from layer 4 while apical dendrites receive feedback, together with three key inhibitory interneuron classes, parvalbumin-expressing (PV), somatostatin-expressing (SOM), and vasoactive-intestinal-peptide-expressing (VIP), that provide compartment-specific inhibition and disinhibition. When input and prediction match, excitation and inhibition remain tightly balanced and activity is sparse; when they differ, this balance is transiently broken in the appropriate compartment, and distinct populations signal either a positive error (unexpected presence) or a negative error (unexpected absence). The circuit reproduces several *in vivo* observations in layer 2/3, including sparse mismatch responses, compartment-specific inhibition, and VIP-mediated disinhibition, and it does so without requiring synaptic weight changes. By routing feature-selective signals from layer 4 into layer 2/3 and relaying the resulting errors toward deeper layers, the model positions layer 2/3 as a local, feature-specific mismatch detector in a hierarchical system. This work provides a concrete, testable mechanism linking dendritic computation, inhibitory diversity, and predictive coding in the cortex.

## 1 Introduction

Predictive coding describes the sensory processing carried out by the cortex as a hierarchical inference engine that continuously compares incoming sensory signals with internally generated expectations [1, 2]. Discrepancies between these two sources of information, referred to as prediction errors (PEs), are believed to update internal models and refine future predictions. Depending on the sign of the mismatch, prediction errors can be positive (PE^+^) when a feature is present in the feedforward input but absent in feedback (FF *>* FB), or negative (PE^−^) when a feature is expected in feedback but missing from the feedforward drive (FB *>* FF) [3, 4]. Identifying the neural circuits that compute PEs is therefore central to understanding how predictive coding could be implemented in the cortical circuit.

Predictive coding has been proposed as a general computational principle of cortical processing, extending beyond the visual system to other sensory and associative areas. Here, we focus specifically on its implementation in the primary visual cortex (V1), where a growing body of experimental evidence localizes prediction-error (PE) signaling to the superficial layers. In behaving mice, two-photon imaging has identified robust and experience-dependent mismatch responses in Layer 2/3 during visuomotor coupling [5, 6]. These signals (i) emerge only after sensorimotor learning [7], (ii) reflect omissions of expected stimuli [8], and (iii) are amplified when somatostatin-expressing (SOM) interneurons are inactivated [9]. Multielectrode array recordings further demonstrate that mismatch-related activity peaks in superficial layers, whereas stimulus adaptation dominates deeper layers [10]. Complementary laminar functional magnetic resonance imaging (fMRI) in humans shows that unexpected stimuli are encoded primarily from superficial voxels, whereas expected stimuli engage the full cortical column [11]. Together, these findings identify Layer 2/3 as a principal locus of visual prediction-error signaling (Table 1).

**Table 1.** Key experimental evidence for prediction errors in visual Layer 2/3.

| Reference | Key Finding | Species | Method |
| --- | --- | --- | --- |
| Keller et al. (2012) [5] | Layer 2/3 neurons respond to visuomotor mismatch; response amplitude scales with running speed | Mouse | 2P $\text{Ca}^{2+}$ |
| Zmarz & Keller (2016) [6] | Layer 2/3 neurons form retinotopically localized mismatch receptive fields during visuomotor coupling | Mouse | 2P $\text{Ca}^{2+}$ (VR) |
| Fiser et al. (2016) [8] | Omission of expected stimuli elicits prediction-error responses after visual experience | Mouse | 2P $\text{Ca}^{2+}$ |
| Hamm et al. (2016) [9] | Prediction-error responses abolished when somatostatin interneurons are silenced | Mouse | LFP + 2P $\text{Ca}^{2+}$ |
| Attinger et al. (2017) [7] | Emergence of prediction-error responses depends on normal visuomotor coupling during development | Mouse | 2P $\text{Ca}^{2+}$ (VR) |
| Jordan & Keller (2020) [12] | Layer 2/3 neurons integrate bottom-up visual and top-down motor inputs, while layer 5 output is suppressed during movement | Mouse | Patch clamp |
| Hamm & Yuste (2021) [10] | Prediction-error activity is strongest in superficial layers, whereas deeper layers show stimulus adaptation | Mouse | Neuropixels |
| Thomas et al. (2024) [11] | Prediction errors decoded primarily from superficial voxels in human V1, while expected stimuli engage the full cortical column | Human | 7T laminar-fMRI |

Building on these empirical findings, computational models of predictive coding have sought to identify how such error signals could emerge from local circuit dynamics. Classical formulations represent PEs as firing-rate variables without explicit cellular or laminar implementation, exchanged between hierarchical layers of the network [1]. Bastos *et al.* mapped this computation onto laminar cortical anatomy while retaining a rate-based formulation [13]. While rate-based models capture predictive coding at a population level, Boerlin *et al.* showed that similar computations can emerge in a spiking network operating in a tightly balanced excitation–inhibition regime, where prediction errors are represented directly in neuronal membrane potentials [14]. This formulation offers a mechanistic bridge between rate models and biophysical cortical circuits, though without explicit feedback pathways.

More recent models have enhanced the biological realism of predictive-coding frameworks along three main dimensions. First, Keller and Mrsic-Flogel proposed distinct excitatory populations for positive and negative prediction errors (PE^+^ and PE^−^), potentially gated by specific interneuron types [3]. Second, dendritic predictive-coding models introduced local comparisons of bottom-up and top-down inputs within pyramidal neurons, showing that basal and apical dendrites can locally compute errors through voltage-dependent coupling [15, 16]. Third, Hertäg and colleagues demonstrated that networks incorporating parvalbumin- (PV), somatostatin- (SOM), and vasoactive-intestinal-peptide- (VIP) expressing interneurons can, through inhibitory homeostatic plasticity, establish stable co-existent PE^+^ and PE^−^ neuronal populations by balancing excitation and inhibition across mismatched input pathways [4]. Subsequently, they extended this framework to encode hierarchical uncertainty, meaning the network estimates and uses precision (inverse variance) of sensory and predictive signals across levels to weight bottom-up vs top-down input streams [17]. Table 2 compares these major models across biological and computational features.

**Table 2.** Computational models of cortical prediction errors.

| Reference | Model Focus / Key Feature | Spiking | 2-Compartment | E-I Balance | Bidirectional PE | Laminar | PV/SOM/VIP | Feedback | Vision |
| --- | --- | --- | --- | --- | --- | --- | --- | --- | --- |
| Rao & Ballard (1999) [1] | Hierarchical rate-based predictive coding with feedforward and feedback pathways | × | × | × | × | × | × | ✓ | ✓ |
| Bastos et al. (2012) [13] | Canonical microcircuit mapping of predictive coding onto cortical laminae | × | × | × | × | ✓ | × | ✓ | × |
| Boerlin et al. (2013) [14] | Balanced spiking network embedding prediction errors in membrane potentials | ✓ | × | ✓ | × | × | × | × | ✓ |
| Keller & Mrsic-Flogel (2018) [3] | Distinct excitatory populations for positive and negative PEs gated by interneurons | × | × | × | ✓ | ✓ | ✓ | ✓ | × |
| Sacramento et al. (2018) [15] | Dendritic predictive coding via local comparison of apical and basal inputs | × | ✓ | ✓ | × | × | ✓ | ✓ | ✓ |
| Mikulasch et al. (2022) [16] | Dendritic error computation in hierarchical cortical inference networks | × | ✓ | ✓ | × | ✓ | ✓ | ✓ | × |
| Hertäg et al. (2022/2025) [4] | PV/SOM/VIP-mediated stabilization of $PE^+$/$PE^-$ populations and uncertainty coding | × | ✓ | ✓ | ✓ | ✓ | ✓ | ✓ | × |
| **This study** | Two-compartment spiking network integrating L4 drive and inhibitory diversity for bidirectional PE computation | ✓ | ✓ | ✓ | ✓ | ✓ | ✓ | ✓ | ✓ |

Despite significant progress across these frameworks, no existing model simultaneously captures dendritic error segregation, inhibitory subtype diversity, laminar connectivity, and spiking dynamics within a unified predictive-coding architecture. Bridging these levels of abstraction is essential for linking the cellular mechanisms of excitation–inhibition balance, where excitatory and inhibitory synaptic inputs dynamically cancel to stabilize activity while preserving sensitivity, to the systems-level process of inference, in which prediction errors iteratively update internal models of sensory input.

The model presented here addresses this gap by embedding the Hertäg & Clopath inhibitory architecture [4] within a two-compartment spiking neuron model that inherits feature-selective feedforward drive from Layer 4. The resulting circuit (i) computes bidirectional prediction errors (PE^+^ and PE^−^) through local soma–dendrite comparisons, (ii) maintains a tight excitation–inhibition balance under matched input but selectively disrupts this balance during mismatches, thereby generating prediction-error responses, and (iii) reproduces hallmark experimental phenomena, including sparse mismatch responses and VIP-mediated disinhibition. By integrating dendritic segregation, interneuron diversity, and laminar-specific spiking dynamics, this model provides a mechanistic account of how Layer 2/3 can operate as a feature-specific mismatch detector and relay error signals to deeper cortical layers for hierarchical inference.

## 2 Model and Methods

The Layer 2/3 (L2/3) circuit was implemented to test how prediction-error (PE) signals arise from realistic local interactions among pyramidal neurons and interneuron subtypes. The model integrates bottom-up feedforward (FF) drive from Layer 4 (L4) with top-down feedback (FB) from higher visual areas (Fig. 1). A detailed description of image preprocessing, parallel ON and OFF pathways from the lateral geniculate nucleus (LGN), which convey increments and decrements in luminance, Gabor-shaped receptive fields modeling the spatial selectivity of thalamocortical afferents, and the Layer 4 spiking network that provides feature-selective feedforward input is provided in Appendices A–D. Local feedforward–feedback comparison produces sign-specific prediction errors: positive (PE^+^) when a feature is present in the feedforward input but absent in feedback (FF *>* FB), and negative (PE^−^) when a feature is expected in feedback but missing from the feedforward drive (FB *>* FF).

**Fig 1.**
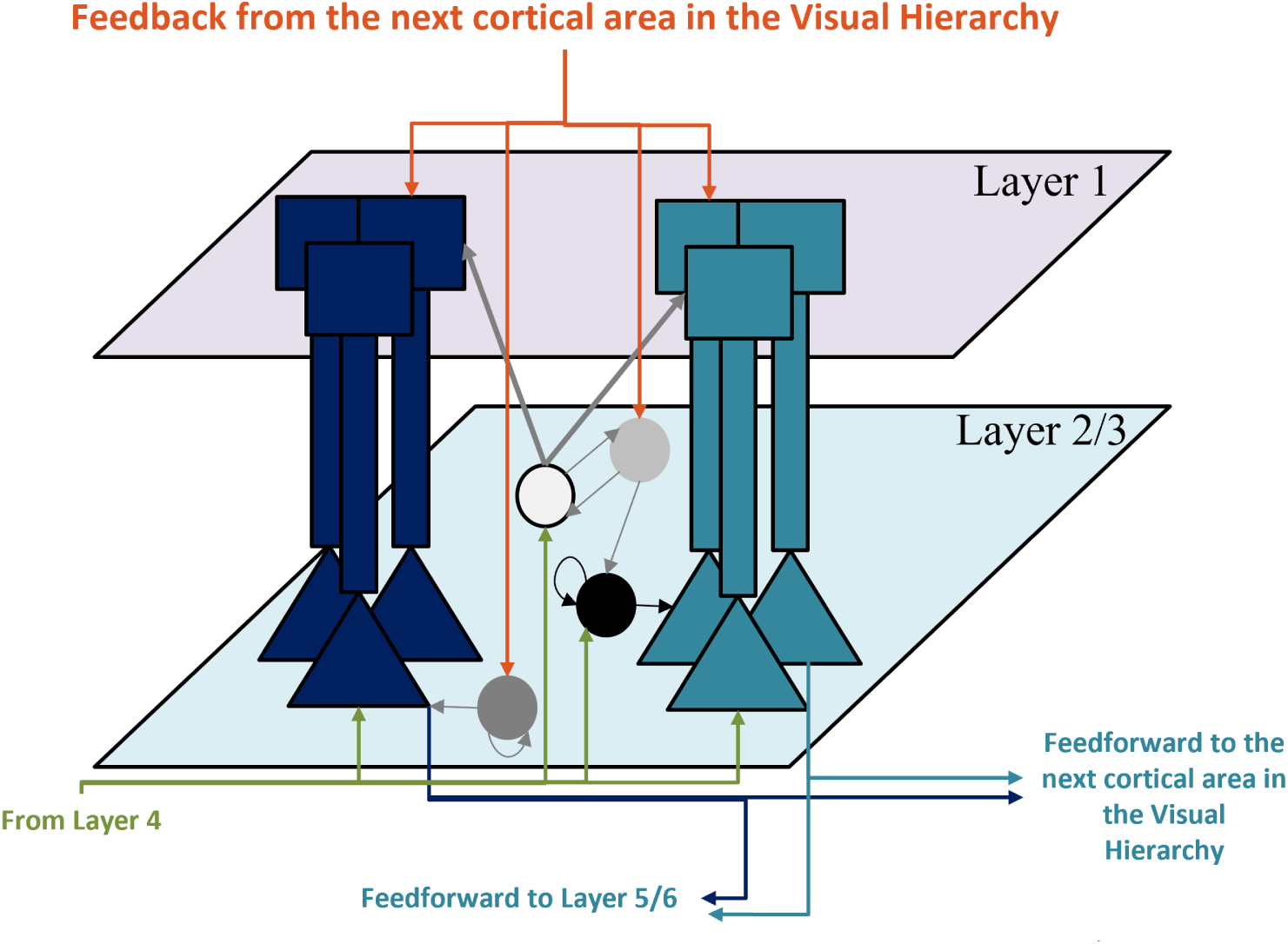
Laminar organisation and information flow in the Layer 2/3 prediction-error circuit. Two-compartment pyramidal neurons encode PE^+^ (navy) and PE^−^ (cyan). Interneurons, parvalbumin (PV, dark grey/black), somatostatin (SOM, white), and vasoactive-intestinal-peptide (VIP, light grey), provide compartment-specific inhibition and disinhibition. Green arrows: feedforward input from Layer 4; orange: feedback to apical tufts and interneurons; navy/cyan: relay of PE signals to deeper layers and other cortical areas; grey: local inhibitory/disinhibitory motifs.

### Populations and roles

Each L2/3 excitatory neuron is modelled as a two-compartment pyramidal cell with distinct somatic and apical integration zones. The somatic compartment receives orientation- and phase-selective feedforward spikes from Layer 4 (feature input), while the apical dendrite receives feedback spikes from higher visual areas via Layer 1 (predictive input). Feature selectivity arises from Gabor-shaped receptive fields that confer tuning to both stimulus orientation and spatial phase, consistent with the thalamocortical input properties of Layer 4 neurons. When feedforward and feedback signals match, excitation and inhibition remain balanced and firing is suppressed; a compartment-specific imbalance activates wither positive (PE^+^) or negative (PE^−^) prediction-error population, corresponding to when the feedforward drive is either stronger or weaker than its feedback prediction, respectively.

Inhibitory control is provided by three major interneuron classes: parvalbumin-expressing (PV) neurons, somatostatin-expressing (SOM) neurons, and vasoactive-intestinal-peptide-expressing (VIP) neurons. PV interneurons are divided into two subpopulations with complementary drives and targets. PV_1_ neurons are driven by feedforward excitation from Layer 4 and inhibit the somata of PE^−^ pyramidal cells, while PV_2_ neurons are driven by feedback excitation and inhibit the somata of PE^+^ cells. This division creates two inhibitory loops, one selective for feedforward-dominated features and the other for feedback-dominated features, allowing each prediction-error population to respond selectively to mismatches in its corresponding input stream [4].

Somatostatin-expressing (SOM) neurons are driven primarily by Layer 4 feedforward input and inhibit both the apical dendrites of pyramidal cells and local VIP neurons when the relevant feature is already represented in the feedforward drive. In this way, SOM cells suppress redundant apical excitation, thereby maintaining dendritic balance when predictions align with sensory input. When a mismatch occurs, apical dendrites become active, allowing excess apical excitation to reach the soma and generate a prediction-error response.

Vasoactive-intestinal-peptide-expressing (VIP) neurons, in turn, receive excitatory feedback drive from higher cortical areas and inhibit SOM neurons. When feedback continues to represent features that are no longer present in the feedforward input, i.e., outdated or mismatched predictions, VIP neurons become active, suppressing SOM inhibition.

Vasoactive-intestinal-peptide-expressing (VIP) neurons receive excitatory feedback input from higher cortical areas. When this feedback continues to predict features that are no longer present in the current feedforward input, VIP neurons become active. Under these mismatch conditions, reduced SOM inhibition allows VIP neurons to inhibit both SOM and PV_2_ interneurons. This dual inhibition transiently releases the apical and somatic compartments of pyramidal cells from inhibition, allowing feedback-driven excitation to activate the corresponding prediction-error population and signal a deviation between prediction and sensory input.

In summary, PV_1_ and PV_2_ interneurons regulate somatic pyramidal activity by suppressing the influence of feedforward or feedback excitation, respectively. SOM neurons suppress apical input when the corresponding feature is already represented in the feedforward stream, while VIP neurons selectively inhibit PV_2_ and SOM interneurons to lift feedback inhibition and permit mismatched predictions to drive the appropriate prediction-error response.

### Circuit scale and constraints

The circuit architecture follows the Layer 2/3 prediction-error microcircuit described by Hertäg et al. (2022) [4], in which excitatory pyramidal neurons receive actual (feedforward) sensory input at the soma and predicted input at the apical compartment, while three inhibitory interneuron classes jointly stabilize network activity through excitation–inhibition (E/I) balance. In our implementation, E/I balance refers to the overall balance between total excitatory and inhibitory input within each compartment, independent of the specific sources of excitation or inhibition. Both populations of PV interneurons provide perisomatic inhibition: PV_1_ neurons are driven by feedforward input and preferentially suppress PE^−^ pyramidal cells, whereas PV_2_ neurons are driven by feedback input and preferentially suppress PE^+^ pyramidal cells, as proposed by Hertäg et al. (2022) [4]. This division of PV neurons into feedforward- and feedback-driven sub-populations is supported by the broad diversity of excitatory inputs to PV interneurons reported experimentally in neocortex [18–21]. SOM interneurons receive strong feedforward excitation from Layer 4 pathways [22], and inhibit both the apical dendrites of pyramidal neurons and local VIP cells [23, 24]. Through this arrangement, SOM neurons maintain dendritic balance when sensory input is accurately predicted. VIP interneurons, in turn, receive excitatory long-range inputs, including top-down feedback from higher cortical areas, and inhibit both SOM and PV_2_ neurons, transiently releasing pyramidal compartments from inhibition when predictions fail to match sensory input [25, 26]. This configuration captures the compartmental excitation–inhibition balance and interneuron connectivity motifs described in mouse V1 [27–29], allowing both PE^+^ and PE^−^ neurons to coexist within the same Layer 2/3 network. A total of 5,632 neurons were simulated within a current-based spiking network model comprising both excitatory and inhibitory cell types. Excitatory neurons were implemented as two-compartment leaky integrate-and-fire (LIF) pyramidal cells representing somatic and apical dendritic compartments, while all interneurons were modeled as single-compartment leaky integrate-and-fire (LIF) units with empirically constrained connectivity patterns (PV→soma; SOM→dendrite, VIP; VIP→SOM, PV_2_) based on mouse V1 microcircuit data [4, 27, 29].

Feedforward synaptic weights from Layer 4 to Layer 2/3 were defined by Gabor filters representing orientation- and phase-selective receptive fields (Appendix B). Feature-aligned connectivity was imposed both within Layer 2/3 (lateral recurrent projections) and between Layer 2/3 and Layer 1 (feedback pathways) by defining synaptic weights as the outer product of pre- and postsynaptic Gabor filters. Following Boerlin et al. (2013) [14], this formulation strengthens the coupling between neurons with similar preferred orientation and spatial phase, maintaining the functional alignment of excitatory and inhibitory populations across laminae. Excitatory-to-excitatory and excitatory-to-inhibitory projections originating from feature-tuned populations followed this rule, while interneurons without direct feedforward input (PV_1_, PV_2_, and VIP) inherited their alignment via excitatory drive. Such spatially organized recurrent and feedback connectivity is consistent with orientation-specific correlations observed experimentally in superficial layers of mouse V1 [30–32].

Interneuron projections were constrained by experimentally established motifs. PV neurons target the somata and proximal dendrites of pyramidal cells [33, 34]; SOM neurons inhibit pyramidal apical dendrites and local VIP cells [22–24, 35]; and VIP neurons inhibit both SOM and subsets of PV interneurons [25, 26].

These connectivity principles were implemented to realize the circuit logic described earlier: PV_1_ and PV_2_ provided somatic inhibition to PE^−^ and PE^+^ populations, respectively; SOM maintained dendritic balance under matched input; and VIP transiently relieved SOM and PV_2_ inhibition during mismatch, producing compartment-specific disinhibition. Functionally, the circuit operates around a dynamic excitation–inhibition (E/I) balance in which inhibitory input conveys the prediction that cancels the excitatory drive when they match. Under this balanced condition, excitatory and inhibitory synaptic currents counteract one another across the soma and dendrites, stabilizing the membrane potential near the resting level and preventing spiking activity. When this balance is perturbed, because sensory excitation diverges from predicted inhibition, the membrane potential transiently depolarizes, generating spikes that decode and propagate the signed prediction error. Thus, correct prediction corresponds to balance, and mismatch corresponds to controlled deviation from balance, consistent with balanced spiking formulations of predictive coding [4, 14].

### Interneuron dynamics

Each interneuron is described by a single-compartment leaky integrate-and-fire (LIF) model. The membrane potential *V_n_*(*t*) evolves according to

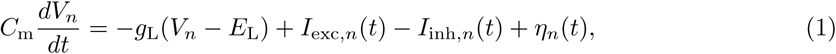

where *C*_m_, *g*_L_, and *E*_L_ are the membrane capacitance, leak conductance, and leak reversal potential. The total input current is the sum of excitatory and inhibitory synaptic currents and a zero-mean Gaussian noise current *η_n_*(*t*) with variance *σ*^2^. Synaptic currents follow first-order kinetics:

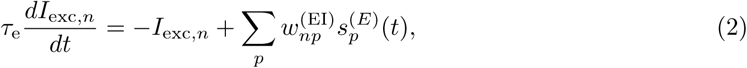

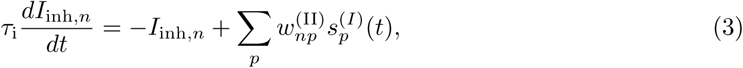

where *I*_exc,*n*_ and *I*_inh,*n*_ denote the total excitatory and inhibitory synaptic currents received by interneuron *n*. Excitatory currents, depending on subtype, originate from feedforward or feedback pathways: PV_1_ and SOM neurons receive feedforward excitation from Layer 4, PV_2_ and VIP neurons receive feedback excitation from higher cortical areas. Inhibitory currents depend on subtype-specific motifs: SOM neurons are inhibited by local VIP cells, VIP neurons inhibit both SOM and subsets of PV interneurons, and PV neurons receive weak mutual inhibition for network stabilization. Each presynaptic spike train *s_p_*(*t*) contributes an instantaneous weight *w_np_* that is exponentially filtered with time constants *τ*_e_ and *τ*_i_, representing excitatory and inhibitory synaptic kinetics.

### Two-compartment pyramidal cell dynamics

Each model pyramidal neuron consists of a somatic and a dendritic compartment coupled through a calcium-dependent current that transmits dendritic activity to the soma. The coupled membrane equations are:

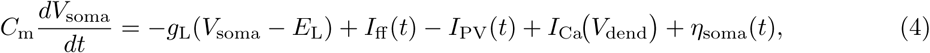

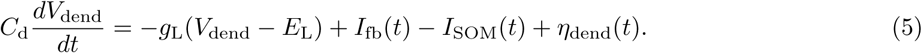

The soma integrates feedforward excitation *I*_ff_ (*t*) opposed by PV-mediated inhibition *I*_PV_(*t*), while receiving a regenerative calcium current *I*_Ca_(*V*_dend_) from the dendrite. The dendritic compartment integrates top-down feedback excitation *I*_fb_(*t*) opposed by SOM-mediated inhibition *I*_SOM_(*t*). Independent Gaussian noise sources *η*_soma_ and *η*_dend_, in the pyramidal neuron soma and dendrite resp., reproduce in vivo voltage fluctuations of 2–5 mV.

When the dendritic potential *V*_dend_ exceeds a calcium threshold (*V*_Ca,th_ = −35 mV), a local plateau current *I*_Ca_ is initiated and sustained for a fixed duration of 50 ms. This current represents a dendritic Ca^2+^ spike that transiently depolarises the soma, enhancing its probability of firing in proportion to dendritic activation. The plateau effectively broadcasts the result of the local dendritic comparison between feedforward and feedback inputs, allowing mismatches in top-down predictions to drive somatic spiking. If the dendritic input remains balanced by SOM inhibition, *V*_dend_ stays below threshold and no plateau is triggered, maintaining quiescence.

PV_1_ and PV_2_ neurons contribute to *I*_PV_(*t*) at the soma, SOM neurons provide dendritic inhibition through *I*_SOM_(*t*), and VIP neurons modulate both by inhibiting SOM and PV_2_. This organisation implements dynamic disinhibition consistent with the microcircuit motifs described above, ensuring that only informative feedforward–feedback mismatches elicit calcium plateau activation and subsequent prediction-error spikes.

### Feedforward input and feature-aligned connectivity

Feedforward drive to Layer 2/3 originates from orientation- and phase-selective excitatory neurons in Layer 4, whose activity is generated through Gabor-based receptive fields and ON/OFF LGN preprocessing (Appendices A–D). Each Layer 4 spike train projects to the somatic compartment of a prediction-error neuron with a matching receptive field, while its apical compartment receives top-down feedback of the same feature class. Synaptic weights are defined as the outer product of pre- and postsynaptic filters [14]:

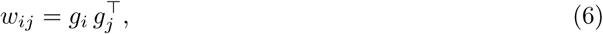

so that neurons with overlapping receptive fields share strong connections, whereas orthogonally tuned pairs interact weakly. The total input current to neuron *i* is

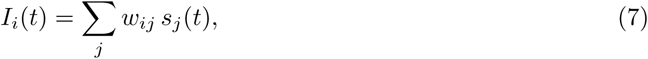

where *s_j_*(*t*) is the presynaptic spike train. Because *w_ij_* inherits the Gabor structure of both neurons, this formulation unifies feedforward, lateral, and feedback inputs within a single feature-aligned basis across soma and dendrite, supporting a coherent computation of prediction errors.

### Parameters

All model parameters were set to match physiological ranges reported for mouse V1 [36]. Excitatory PE^+^ and PE^−^ neurons were implemented as two-compartment pyramidal cells, while PV_1_, PV_2_, SOM, and VIP neurons were single-compartment leaky integrate-and-fire (LIF) units. Synaptic weights were scaled by a global factor *w*_0_ chosen such that a single excitatory synapse evoked a postsynaptic current of approximately 10–20 pA at rest, matching unitary EPSC amplitudes reported in mouse V1 [27]. Each neuron received independent zero-mean Gaussian current noise (RMS ≈20 pA), introducing realistic subthreshold fluctuations (2–5 mV) consistent with in vivo recordings. Feedforward input was provided by the Layer 4 spiking network defined in Appendices A–D, ensuring matched feature statistics and receptive-field alignment across hierarchical levels. Key biophysical and simulation parameters are summarised in Table 3.

**Table 3.** Model parameters for the Layer 2/3 predictive-coding network.

| Parameter | Value (biological range) |
| --- | --- |
| Input patch size | $16 \times 16$ pixels ($\sim 2^\circ$ visual angle) |
| Number of LGN neurons | 256 ON + 256 OFF |
| Number of L4 neurons | 4,096 excitatory, 1,024 inhibitory |
| Number of L2/3 neurons | 2,048 PE<sup>+</sup>, 2,048 PE<sup>-</sup>, 512 PV<sub>1</sub>, 512 PV<sub>2</sub>, 256 SOM, 256 VIP |
| Unit weight scale $w_0$ | 0.5 nS (EPSP $\approx 15$ pA) |
| Simulation time step $\Delta t$ | 0.1 ms |
| Membrane time constant $\tau_m$ | 10 ms (all neuron types) |
| Membrane capacitance $C_m$ | 200 pF (soma), 100 pF (dendrite), 150 pF (interneuron) |
| Leak potential $E_L$ | -70 mV |
| Spike threshold $V_{th}$ | -50 mV |
| Reset potential $V_r$ | -60 mV |
| Refractory period $\tau_r$ | 2 ms (excitatory), 1 ms (inhibitory) |
| Synaptic time constants $\tau_e, \tau_i$ | 5 ms (excitatory), 10 ms (inhibitory) |
| Reversal potentials $E_E, E_I$ | 0 mV, -80 mV |
| Ca<sup>2+</sup> threshold $V_{Ca,th}$ | -35 mV |
| Ca<sup>2+</sup> plateau duration | 50 ms |
| Noise RMS $\sigma$ | 20 pA (2–5 mV fluctuations) |
| Gabor filter size | $16 \times 16$ pixels; aspect ratio $\gamma = 0.5$ |
| Orientation/phase sampling | 10 orientations × 10 phases (uniform) |

### 2.1 Functional Measures and Network Readouts

To evaluate how effectively the Layer 2/3 microcircuit encodes prediction errors, we quantified (i) the selectivity of population firing responses under canonical feedforward–feedback (FF–FB) configurations, and (ii) compartment-specific excitation–inhibition (E–I) balance within somatic and dendritic compartments. All measures were computed from steady-state spike trains and filtered synaptic currents (10 ms exponential kernel) recorded during the four canonical conditions: positive error (FF present, FB absent), negative error (FB present, FF absent), mismatch (both present but differing), and full prediction (identical FF and FB).

#### Prediction–error selectivity

For each neuronal population (PE^+^, PE^−^, PV_1_, PV_2_, SOM, VIP), we measured the mean firing rate under each condition and defined a normalized selectivity index:

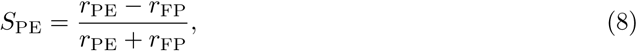

where *r*_PE_ and *r*_FP_ denote the mean firing rates during the preferred prediction-error and fully predicted (baseline) conditions, respectively. This index ranges from −1 (complete suppression) to +1 (maximal selective activation) and quantifies each population’s ability to signal unexpected sensory events while remaining silent under accurate predictions.

#### Excitation–inhibition balance

To characterise the underlying circuit dynamics, somatic and dendritic current traces were analysed using three current-based measures following Ahmadian & Miller (2021) [37]. Excitatory (*I*_exc_) and inhibitory (*I*_inh_) currents were smoothed (10 ms window) and averaged over trials.

##### 1 Linear slope

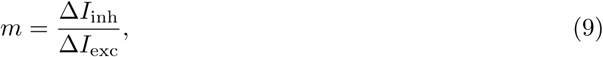

estimated by linear regression of inhibitory on excitatory current amplitudes. A slope near 1 indicates proportional coupling between excitation and inhibition.

##### 2 Correlation coefficient

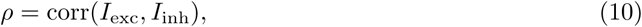

which quantifies the temporal co-fluctuation of excitatory and inhibitory inputs; higher *ρ* values denote tighter E–I synchrony.

##### 3 Balance index

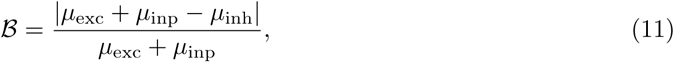

where *µ*_exc_, *µ*_inh_, and *µ*_inp_ are the mean excitatory, inhibitory, and external currents. This index captures the deviation of total synaptic input from perfect cancellation: B ≪ 0.1 denotes tight balance, whereas larger values indicate a transient imbalance that enables spiking output.

Together, these measures provide a quantitative measure of how local interneuron interactions regulate compartmental E–I coordination and thereby determine the precision and sign specificity of prediction–error signalling in Layer 2/3.

## 3 Results

This study presents a biophysically realistic spiking model of Layer 2/3 in V1 to investigate how local cortical microcircuits generate and encode prediction errors. Each pyramidal neuron was represented by two compartments: a somatic compartment receiving feedforward input from Layer 4, and an apical dendritic compartment receiving feedback input from higher visual areas. Three classes of inhibitory interneurons, parvalbumin-expressing (PV), somatostatin-expressing (SOM), and vasoactive-intestinal-peptide-expressing (VIP), provided pathway-specific inhibition and disinhibition to regulate compartmental activity.

The circuit was tested under four canonical combinations of feedforward and feedback stimulation, corresponding to distinct sensory–predictive relationships as defined in the Model and Methods Section 2.

The analysis proceeded in three stages, each addressing a distinct aspect of prediction-error computation in the model. First, we established the expected pattern of somatic and dendritic activation across the four canonical feedforward–feedback conditions and outlined the inhibitory pathways that shape these responses (see Figure 2 and Section 3.1). This step defined the mechanistic logic of the circuit, namely how excitation, inhibition, and disinhibition should interact under positive, negative, matched, and mismatched input configurations.

**Fig 2.**
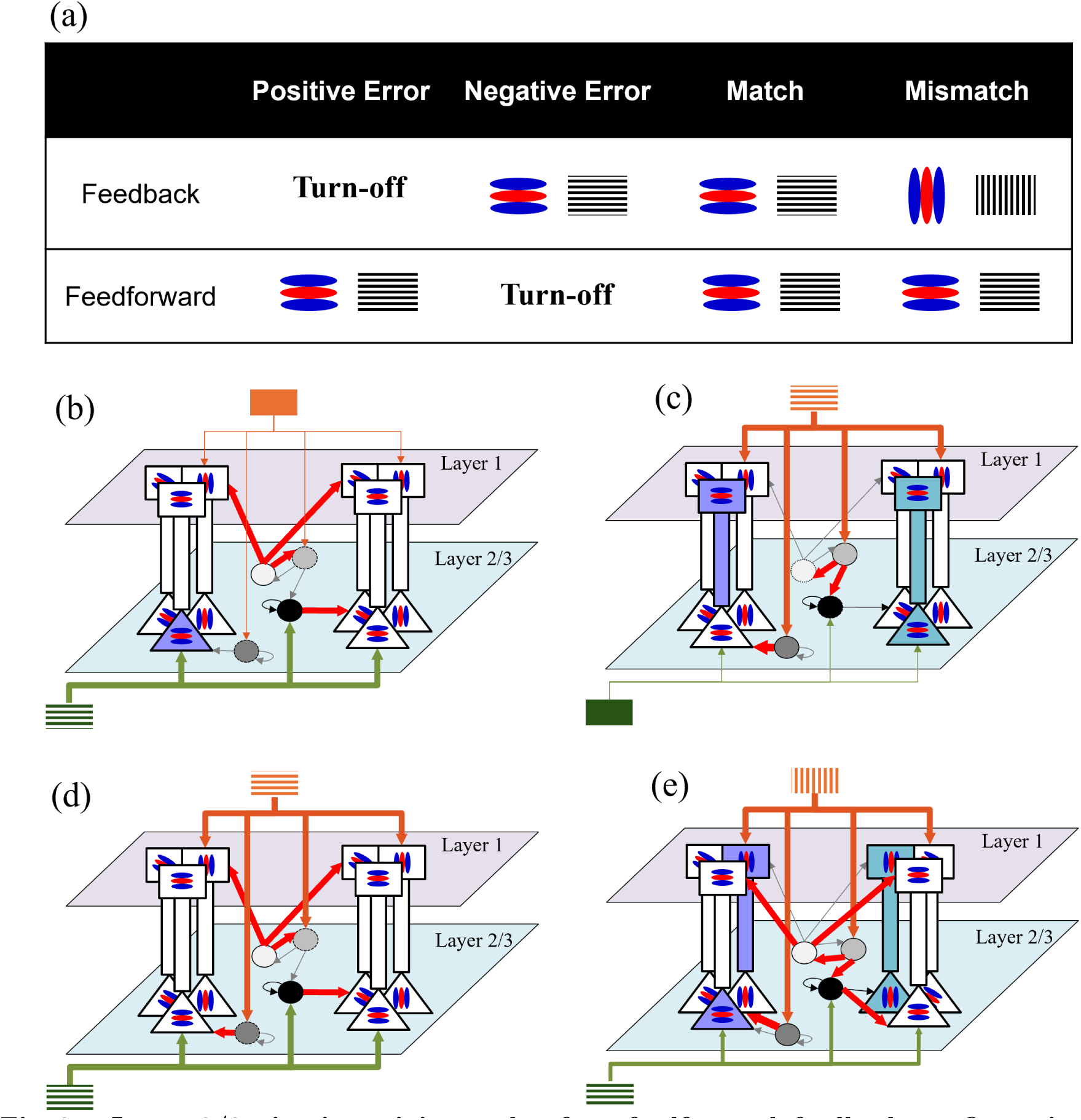
Layer 2/3 circuit activity under four feedforward–feedback configurations. (a) Canonical stimulus conditions: *Positive prediction error*, PE^+^ (feedback absent), *Negative prediction error*, PE^−^ (feedforward absent), *Match* (feedforward equals feedback), and *Mismatch* (feedforward and feedback differ in feature identity). (b–e) Schematic pathway diagrams for each condition. Two-compartment pyramidal neurons encode positive (navy) and negative (teal) prediction errors by comparing somatic and dendritic inputs. Interneurons, PV_1_, PV_2_, SOM, and VIP, provide compartment-specific inhibition and disinhibition. Filled symbols denote active populations; thick red arrows indicate those pathways engaged in each scenario.

Second, we analysed the firing activity of somatic and dendritic compartments (Section 3.2) to test whether the excitatory populations representing positive and negative prediction errors respond selectively to their respective mismatch type. Somatic activity was expected to report the unexpected appearance of a sensory/feedback feature in the feedforward input (positive/negative prediction error), while dendritic activity was expected to signal a new or unconfirmed feature in the feedback signal that was not supported by the sensory evidence. To quantify these effects, three complementary excitation–inhibition metrics were computed: (i) the linear excitation–inhibition slope, (ii) the excitatory–inhibitory correlation coefficient, and (iii) the balance index (as defined in Section 2.1), to examine how synaptic balance shifts during prediction and mismatch conditions. The results are shown in Figures 3 and 4.

**Fig 3.**
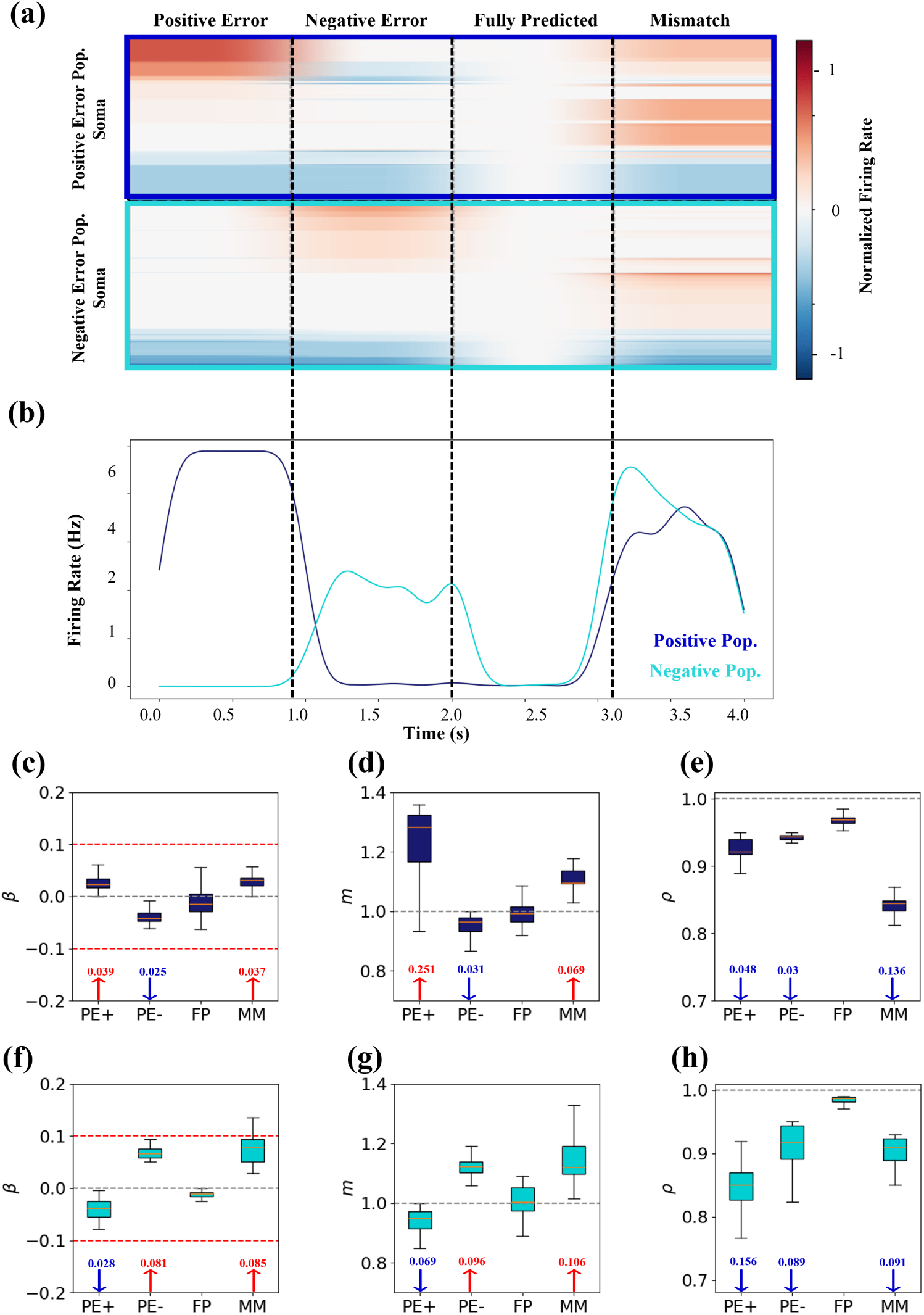
Somatic activity and balance metrics of prediction-error neurons. **(a)** Normalized firing-rate heatmaps for PE^+^ (top) and PE^−^ (bottom) somata, computed relative to each neuron’s 500 ms pre-stimulus baseline and scaled to its maximum absolute deviation across all conditions. **(b)** Population-averaged firing rates (mean ± SEM) within each 1 s stimulus window. **(c–e)** E–I balance index *β*, excitation–inhibition slope *m*, and correlation *ρ* for PE^+^ neurons. **(f–h)** Same metrics for PE^−^ neurons. Red (blue) arrows indicate increases (decreases) relative to the fully predicted (FP) baseline; dashed red lines mark the tight-balance band |*β*| *<* 0.1.

**Fig 4.**
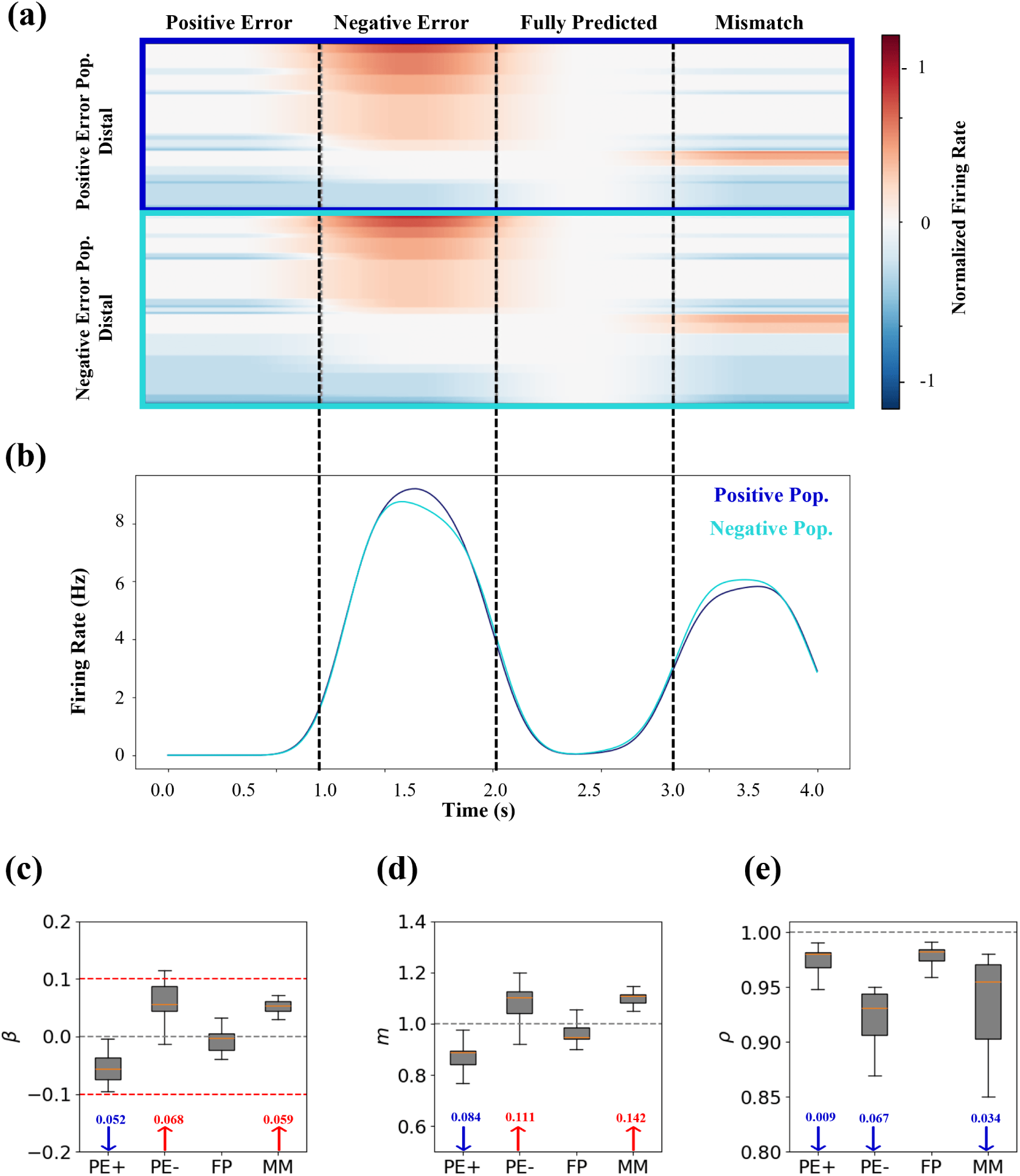
Distal compartment activity and excitation–inhibition (E–I) coordination metrics. **(a)** Normalized dendritic firing-rate heatmaps computed relative to each neuron’s 500 ms pre-stimulus baseline and scaled to its maximum absolute deviation across all conditions. **(b)** Population-averaged firing rates (mean ± SEM) within each 1 s stimulus window, showing strong activation during PE^−^ and mismatch. **(c–e)** E–I balance index *β*, E–I input slope *m*, and excitatory–inhibitory correlation *ρ*, computed relative to the fully predicted (FP) baseline. Red (blue) arrows indicate increases (decreases) relative to FP; dashed red lines mark the tight-balance band |*β*| *<* 0.1.

Finally, we extended the same analyses to the inhibitory populations to assess how parvalbumin- (PV), somatostatin- (SOM), and vasoactive-intestinal-peptide-expressing (VIP) interneurons regulate excitation–inhibition coordination across conditions. These interneuron classes were expected to maintain a tight balance during accurate predictions and transiently release inhibition when mismatches occur. The same three quantitative measures, the linear slope, correlation coefficient, and balance index, were measured to demonstrate how each population dynamically gates pyramidal activity. The results, illustrated in Figures 5 and 6, show that inhibitory populations preserve near-perfect balance under matched input while modulating inhibition selectively in response to prediction errors.

**Fig 5.**
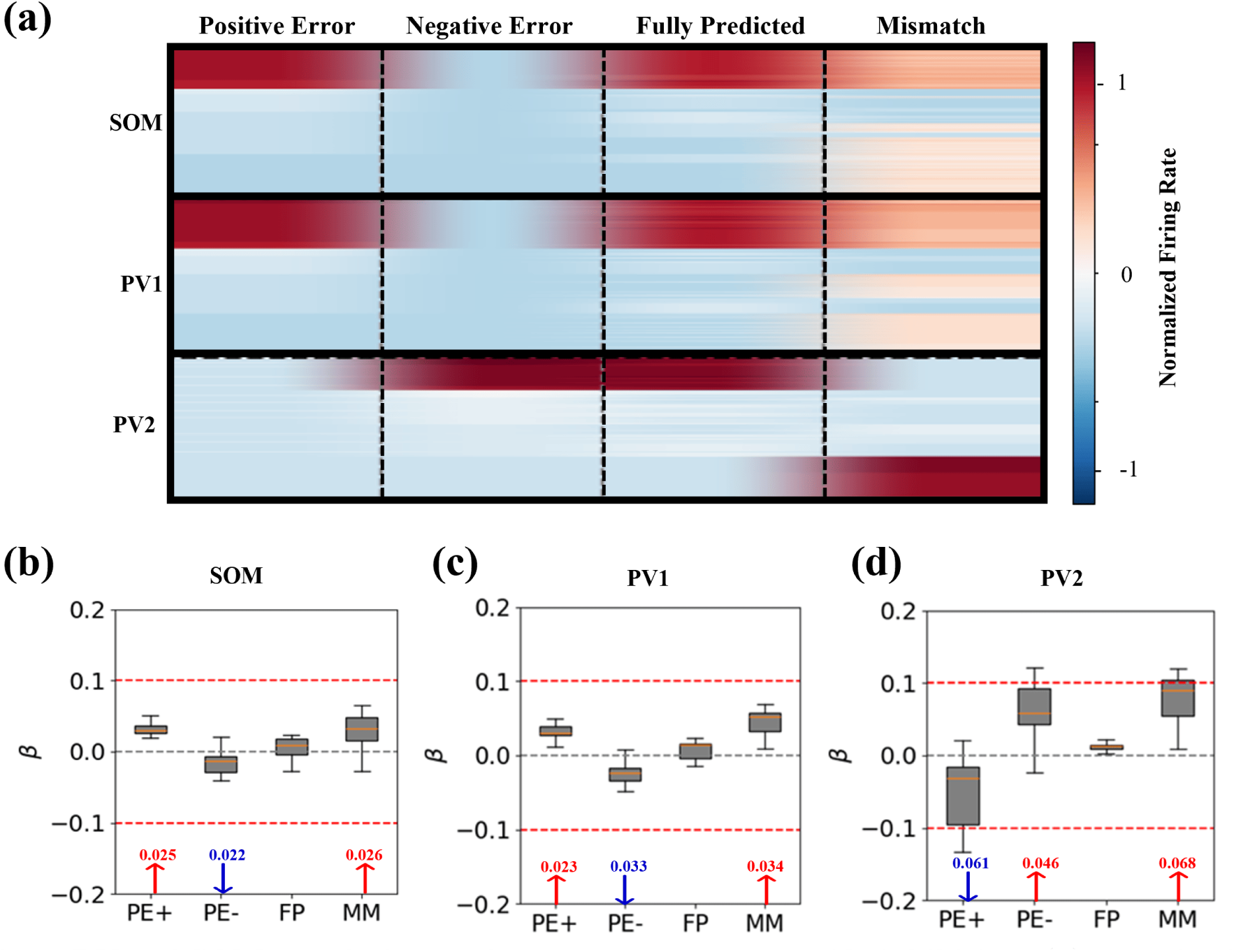
PV and SOM populations enforce compartment-specific inhibition. **(a)** Normalized firing-rate heatmaps for SOM, PV_1_, and PV_2_, computed relative to each neuron’s 500 ms prestimulus baseline. **(b–d)** E–I balance index *β* for each population across conditions. Red (blue) labels denote signed deviations from the fully predicted (FP) baseline. All distributions remain within the tight-balance range (|*β*| *<* 0.1).

**Fig 6.**
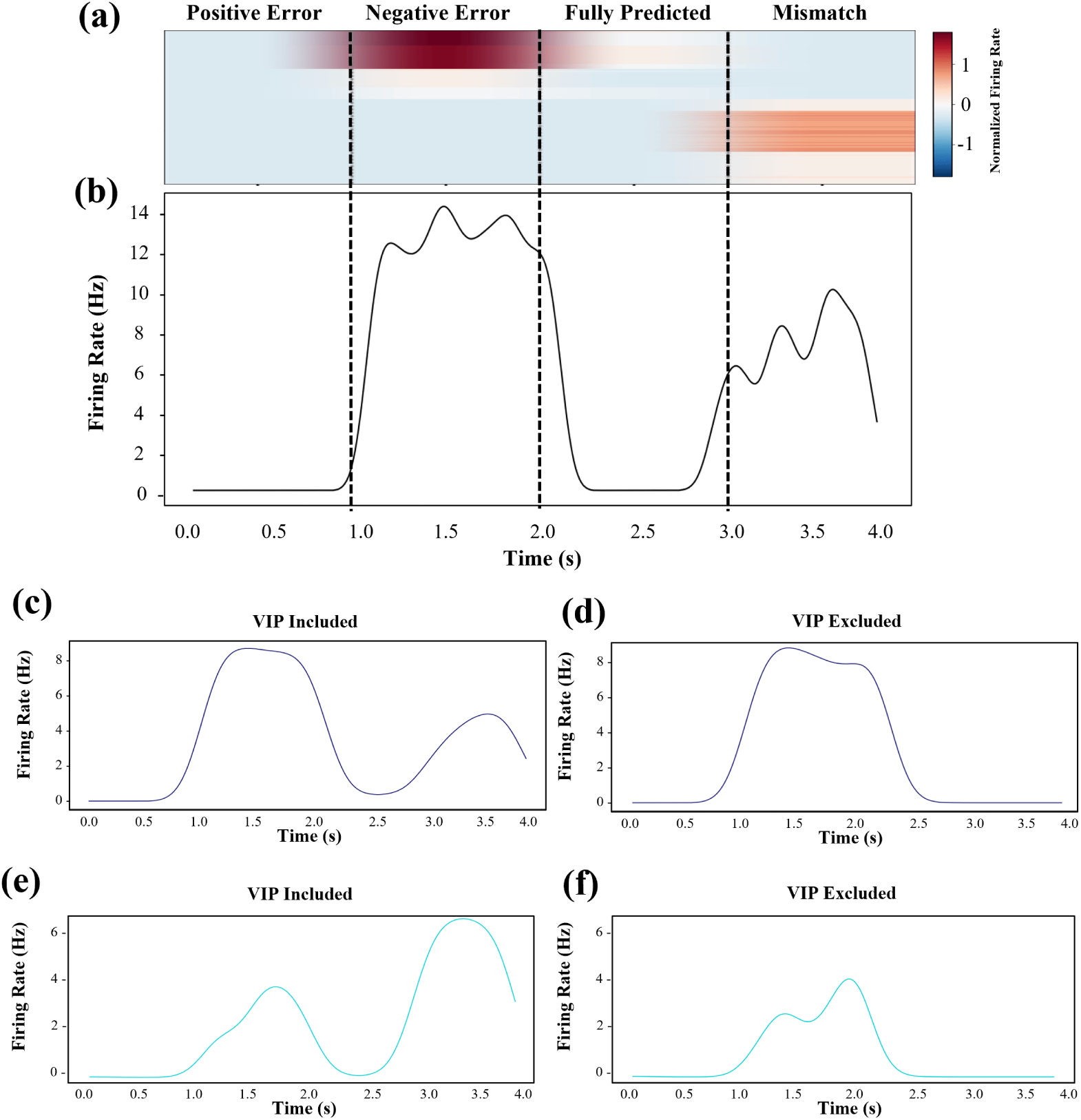
VIP neurons mediate feedback-dependent disinhibition and gating of prediction errors. **(a)** Normalized firing-rate heatmap for VIP neurons across the four canonical conditions, showing selective activation during feedback-present conditions (PE^−^ and mismatch, MM). **(b)** Population-averaged VIP firing rate over time, peaking transiently during feedback-driven mismatch. **(c–d)** Dendritic prediction-error (PE) activity with (c) and without (d) VIP output, demonstrating loss of mismatch responses when disinhibition is removed. **(e–f)** Somatic PE^−^ activity with (e) and without (f) VIP output, confirming that VIP-mediated disinhibition is required for feedback-related PE signaling.

The results reveal a clear and consistent pattern. Neurons representing positive prediction errors, PE^+^, fire selectively when an unexpected sensory feature is present, whereas neurons representing negative prediction errors, PE^−^, fire when a predicted feature is missing. When feedforward and feedback inputs match, both populations remain near baseline, indicating accurate prediction. In all conditions, non-engaged compartments maintain a tight excitation–inhibition balance, demonstrating that pathway-specific inhibitory control supports a sparse and sign-specific representation of prediction errors in Layer 2/3.

### 3.1 Prediction–error scenarios and expected circuit responses

To evaluate how the Layer 2/3 microcircuit encodes prediction errors, we simulated four canonical combinations of feedforward and feedback input, targeting the somatic and apical compartments independently. Each configuration represents a distinct sensory–predictive relationship and activates a specific pattern of excitatory and inhibitory pathways within the circuit.

#### Positive prediction error, PE^+^ (feedback absent)

Only feedforward drive reaches the soma, representing an unexpected sensory feature. PV_1_ and SOM neurons, both driven by feedforward input, suppress the somata of negative-error cells and all apical dendrites, preventing spurious feedback activation. PV_2_ and VIP remain silent. As a result, positive-error neurons fire strongly, signalling the unexpected presence of a feature in the sensory input.

#### Negative prediction error, PE^−^ (feedforward absent)

Only feedback reaches the apical dendrites, representing a predicted feature that fails to appear in the sensory input. The resulting dendritic depolarization generates a calcium plateau potential that drives somatic spiking in negative-error neurons. Feedback-driven PV_2_ interneurons inhibit the somata of positive-error neurons, while PV_1_ and SOM remain inactive.

#### Match (feedforward equals feedback)

When sensory input and feedback prediction are identical, both pathways are co-active. PV_1_, PV_2_, and SOM interneurons are jointly recruited, clamping somatic and dendritic compartments in tight balance. VIP cells remain silent, and no prediction-error signal is generated, reflecting a state of accurate prediction.

#### Mismatch (feedforward and feedback differ)

Feedforward input excites one set of positive-error somata, while feedback excites a separate set of negative-error dendrites tuned to different features. Because each pathway engages the other population’s inhibitory partners (PV and SOM), mis-matched compartments temporarily escape suppression, resulting in simultaneous, feature-specific activation of both positive- and negative-error neurons. VIP neurons are transiently activated by the feedback component, releasing inhibition and amplifying the mismatch response.

Together, these four scenarios illustrate how compartment-specific inhibition and dendritic non-linearity enable the Layer 2/3 circuit to encode the full spectrum of sensory–predictive relations: the unexpected presence of a feature (positive error), the unexpected absence of a feature (negative error), a perfect match (accurate prediction), and a partial conflict between the two streams (mismatch).

### 3.2 Activity of Prediction-Error Neurons

To examine how excitatory neurons encode prediction errors, we analysed the somatic and dendritic activity of *N*_PE_+ = 998 positive-error and *N*_PE_*−* = 1023 negative-error neurons that exceeded the five-spike inclusion threshold. These cells were drawn from a Layer 2/3 network of 4096 two-compartment pyramidal neurons (2048 of each type). Responses were evaluated across four stimulus configurations corresponding to positive error (PE^+^), negative error (PE^−^), fully predicted (FP), and mismatch (MM) conditions (Figure 3).

#### Somatic firing and balance

Figure 3a shows normalized somatic firing rates for both prediction-error populations, scaled from −1 (suppressed) to +1 (maximally active) relative to each neuron’s own baseline. Normalization was performed by subtracting the mean firing rate in a 500 ms pre-stimulus window and dividing by the maximum absolute deviation across all stimulus conditions, yielding a dynamic range from −1 to +1 per neuron. In this scale, +1 indicates peak excitation, 0 represents baseline, and −1 corresponds to complete suppression.

During the PE^+^ condition, positive-error neurons reached near-maximal activation while negative-error neurons were strongly inhibited; this pattern reversed in the PE^−^ condition. Both populations remained near baseline in the fully predicted (FP) condition and fired moderately during mismatch (MM), reflecting the concurrent feedforward and feedback drive.

Mean firing rates (Figure 3b) confirmed these selectivities: PE^+^ neurons peaked at 7.2 ± 0.6 Hz during PE^+^, PE^−^ neurons at 2.6 ± 0.8 Hz during PE^−^, and both at approximately 5 Hz under mismatch (MM), remaining close to 1 Hz in the fully predicted (FP) condition.

Excitation–inhibition coordination was quantified within each 1 s stimulus window using three complementary metrics (Figures 3c–h): the normalized balance index *β*, the linear slope *m* between inhibitory and excitatory currents, and the Pearson correlation coefficient *ρ*. For each neuron, the baseline reference was taken from the fully predicted (FP) condition, and all reported changes (Δ) indicate deviations from this reference value.

In PE^+^ neurons, excitation dominated during the PE^+^ window (Δ*β* = +0.039, Δ*m* = +0.25, Δ*ρ* = −0.05), indicating a transient shift toward stronger excitation and weaker input coupling. A smaller imbalance persisted during mismatch (Δ*m* = +0.07). The inverse pattern was observed in PE^−^ neurons during the PE^−^ window (Δ*β* = +0.05, Δ*m* = +0.10, Δ*ρ* = −0.09), consistent with enhanced excitatory drive under feedback mismatch (MM). In the FP condition, both populations maintained tight balance (|*β*| *<* 0.1, *ρ >* 0.95), confirming that inhibition precisely tracks excitation during accurate prediction.

#### Dendritic firing and balance

Dendritic activity (Figure 4) captured the complementary, feedback-driven component of the prediction-error code. Normalization was performed per neuron by subtracting its 500 ms pre-stimulus baseline and dividing by the maximum absolute deviation across all stimulus conditions, yielding values from −1 (suppressed) to +1 (maximally active). Within each 1 s stimulus window, distal compartments showed strong activation during the PE^−^ and mismatch (MM) conditions and remained near baseline during PE^+^, confirming selectivity for feedback mismatch. Average firing rates reached 8.6 ± 0.7 Hz in PE^−^ and 6.4 ± 0.6 Hz in MM, both significantly above the fully predicted baseline.

Excitation–inhibition coordination followed the same pattern (Figures 4c–e). All reported deviations (Δ) are relative to the FP baseline. During PE^−^, the balance index increased by Δ*β* = +0.07, the E–I input slope by Δ*m* = +0.11, and the E–I correlation declined by Δ*ρ* = −0.07. Similar but slightly weaker shifts were observed during mismatch (Δ*β* = +0.06, Δ*m* = +0.14, Δ*ρ* = −0.03), indicating transient dominance of excitation and weakened inhibitory coupling when feedback predictions diverge from sensory evidence.

Together, these simulation results show that prediction-error neurons operate through compartment specific E–I imbalance. Somatic compartments encode unexpected feedforward/feedback drive by escaping PV-mediated inhibition, whereas dendritic compartments encode unexpected or unconfirmed feedback through release from SOM-mediated inhibition. Under matched input, both compartments remain tightly balanced (|*β*| *<* 0.1, *ρ >* 0.95), preventing spurious error output and preserving metabolic efficiency.

### 3.3 Inhibitory Population Dynamics

We next examined how parvalbumin- (PV), somatostatin- (SOM), and vasoactive-intestinal-peptide- (VIP) expressing interneurons maintain excitation–inhibition (E–I) balance and dynamically gate prediction-error signaling (Figures 5, 6). All analyses were performed within 1 s stimulus windows following a 500 ms pre-stimulus baseline, using the same normalization and E–I metrics as in the excitatory populations.

#### PV and SOM: compartment-specific inhibition

Figure 5a shows normalized firing-rate heatmaps for SOM, PV_1_, and PV_2_ neurons. PV_1_ and SOM, both driven by feedforward input, inhibit the somatic and dendritic compartments respectively whenever bottom-up drive is present. In contrast, PV_2_ is recruited by feedback predictions and suppresses somatic output when top-down signals dominate, thereby maintaining compartment-specific inhibitory control.

The E–I balance distributions (Figures 5b–d) confirmed this functional specialization across interneuron types. Relative to the fully predicted (FP) baseline, PV_1_ and SOM exhibited decreased balance values during PE^−^ (Δ*β* ≈ −0.03), while remaining elevated during PE^+^, FP, and mismatch (MM) conditions. PV_2_ showed the opposite pattern, peaking during PE^−^ and MM conditions (Δ*β* ≈ +0.07), reflecting its feedback-driven activity. Although all populations stayed within the tight-balance band (|*β*| *<* 0.1), these coordinated shifts demonstrate how PV and SOM interneurons enforce prediction-dependent inhibition across somatic and dendritic compartments.

#### VIP: feedback-driven disinhibition

VIP neurons act indirectly by targeting SOM and PV_2_, thereby releasing dendritic and somatic inhibition when feedback mismatch occurs. As shown in Figures 6a–b, VIP firing remained minimal during PE^+^ and FP condtions but increased sharply during PE^−^ and MM conditions, coinciding with the onset of feedback input. This activity enables selective disinhibition when top-down predictions conflict with sensory evidence.

Eliminating VIP output abolished this disinhibitory mechanism: dendritic mismatch responses were markedly attenuated, and somatic PE^−^ activity was almost completely suppressed (Figures 6c–f). These findings demonstrate that VIP-mediated disinhibition is a necessary mechanism for enabling feedback-driven prediction-error signaling.

Collectively, the inhibitory populations sustain a near-perfect excitation–inhibition balance during accurately predicted inputs but transiently relax inhibition when mismatches arise. This dynamic regulation permits compartment-specific excitation to emerge selectively during sensory–predictive conflict, thereby supporting a sparse, metabolically efficient representation of prediction errors in Layer 2/3.

## 4 Discussion

This study presented a biologically grounded spiking model of cortical Layer 2/3 that computes prediction errors through compartment-specific interactions among pyramidal neurons and inhibitory interneurons. Building upon the structured sensory representations generated in Layer 4, the model integrates bottom-up sensory drive at the soma with top-down feedback at the apical dendrite of two-compartment pyramidal neurons. By incorporating three inhibitory classes, parvalbumin-expressing (PV), somatostatin-expressing (SOM), and vasoactive-intestinal-peptide-expressing (VIP) cells, the circuit achieves compartment-specific gating of excitation and encodes bidirectional prediction errors, PE^+^ and PE^−^, consistent with predictive coding theory. The following discussion addresses the model’s implications for cortical inference, biological plausibility, generalization across modalities, and emergent coding principles, and outlines potential directions for future work.

### Predictive coding perspective

The model formalizes prediction-error computation as the comparison between feedforward input arriving at the soma and feedback arriving at the apical dendrite. When these inputs are balanced, both compartments remain near baseline, representing an accurate prediction. When balance is broken, distinct populations of excitatory neurons signal positive or negative errors, depending on whether the mismatch arises from the unexpected presence or absence of sensory features. Somatic activity corresponds to positive/negative errors driven by unpredicted feedforward/feedback input, while dendritic activity reflects feedback predictions unsupported by sensory evidence. This bidirectional error representation aligns with recent formulations of dendritic predictive coding [3, 4]. It provides a mechanistic explanation for how pyramidal neurons can integrate and compare bottom-up and top-down information within their cellular architecture.

By mapping opposite error types onto separate excitatory pyramidal populations, the model supports sign-specific signalling that can be locally computed and hierarchically propagated. Unlike abstract rate-based models [1, 4], prediction errors in this framework emerge from transient deviations in local excitation–inhibition coordination rather than explicit subtraction operations. This architecture enables a sparse, feature-selective, and energetically efficient representation of unexpected events within the canonical cortical microcircuit.

### Biological plausibility

The circuit design adheres to key anatomical and physiological constraints of the visual cortex, supported by extensive physiological data from mouse visual cortex. Two-compartment excitatory neurons reflect the known separation of feedforward and feedback inputs: ascending drive from Layer 4 targets the soma, whereas descending feedback from higher areas targets apical dendrites in Layer 1. The nonlinear coupling between these compartments allows dendritic plateau potentials to trigger somatic spiking only under mismatch conditions, consistent with experimental findings in pyramidal neurons [38]. PV, SOM, and VIP interneurons were implemented as single-compartment spiking units with realistic connectivity patterns. PV neurons mediate perisomatic inhibition and stabilize baseline activity, SOM neurons inhibit distal dendrites to regulate apical input, and VIP neurons provide conditional disinhibition by targeting SOM and PV sub-types. These motifs reproduce experimentally observed patterns of inhibitory control [4, 27] and demonstrate how balanced and disinhibitory mechanisms can jointly implement predictive inference. A central feature of this model is that compartment-specific excitation–inhibition balance is preserved under predicted input but is selectively disrupted during mismatches. Quantitative analyses of the balance index, current correlation, and E–I slope show that prediction errors correspond to brief, structured deviations from balance rather than absolute increases in excitation. This organization explains how cortical circuits can remain globally stable while being locally responsive to prediction violations.

### Generalization across modalities and cortical areas

Although the model was developed for the primary visual cortex, the same computation—comparison of feedforward and feedback signals under interneuron-specific gating, likely generalizes across sensory and associative cortices. The canonical Layer 2/3 motif comprising pyramidal neurons, PV, SOM, and VIP interneurons recurs throughout neocortex, suggesting a universal computational role. In other modalities, the function of the Layer 4 encoder would be implemented by modality-specific feature filters: spectrotemporal receptive fields in auditory cortex, spatiotemporal receptor filters in somatosensory cortex, or complex feature detectors in higher visual areas. The Layer 2/3 microcircuit would then compute prediction errors by comparing sensory and predictive inputs within the same compartment.

The laminar routing of feedforward input through Layer 4 and feedback through Layer 1 is highly conserved, implying that similar positive and negative prediction error signals, PE^+^ and PE^−^, should occur across sensory systems. In the auditory cortex, the omission of an expected tone should activate negative-error neurons, while unexpected tactile input in the somatosensory cortex should activate positive-error neurons. This framework predicts that feedback-driven mismatch responses will depend critically on VIP-mediated disinhibition, a hypothesis testable with optogenetic perturbations or two-photon calcium imaging.

### Interneuron-specific control and emergent coding principles

The inhibitory cortical microcircuit described in this study forms a flexible control system that regulates when and where prediction errors emerge. PV neurons enforce somatic inhibition and prevent spurious firing during predicted input. SOM neurons maintain dendritic balance, limiting apical excitation until feedback mismatch occurs. VIP neurons transiently disinhibit both soma and dendrite during feedback violations, allowing PE^−^ neurons to signal missing or unexpected features. Removing VIP output abolishes these feedback-driven responses, confirming its role as a disinhibitory gate. This dynamic gating reproduces experimentally observed inhibitory interactions in superficial layers and supports the view that cortical balance is dynamically modulated rather than static [39, 40].

Across all conditions, the circuit maintains a tight excitation–inhibition balance during accurate predictions but selectively relaxes this balance during mismatches. The resulting spiking activity is sparse, sign-specific, and compartmentalized, all properties that mirror physiological recordings from Layer 2/3 in behaving animals under visuomotor and omission paradigms [5, 9, 12]. These results suggest that cortical microcircuits implement predictive inference not through global modulation or explicit subtraction, but through precise, transient shifts in local balance controlled by inhibitory cortical microcircuit motifs.

### Broader significance

This framework bridges cellular biophysics with theories of cortical inference by showing how prediction and error emerge from the same cortical microcircuit through different states of balance. Prediction corresponds to equilibrium, while prediction error corresponds to the transient imbalance between feedforward and feedback pathways. Such compartment-specific modulation enables the cortex to remain stable yet adaptive, allowing sparse, energy-efficient coding of unexpected events. By grounding predictive coding in experimentally accessible mechanisms, this work offers a unifying account of how cortical microcircuits transform synaptic dynamics into inference that is experimentally testable.

### 4.1 Future directions

The Layer 2/3 model presented here establishes a biologically plausible circuit for computing direction-specific prediction errors based on local somato-dendritic comparisons. Building on this foundation, several directions can extend the framework toward a complete hierarchical and dynamic model of cortical inference.

#### Link to hierarchical predictive coding in deeper layers

The present model defines Layer 2/3 as the computational locus for generating signed prediction errors that provide the bottom-up teaching signal for deeper cortical layers. In future work, this architecture will be extended to Layer 5/6, which is proposed to generate top-down predictions based on integrated error signals. This addition will enable a closed-loop implementation of predictive coding, in which descending predictions iteratively minimize superficial-layer error activity. Such an extension would allow the entire cortical column to perform dynamic inference, in which prediction errors are progressively suppressed as internal models adapt and converge toward the sensory input.

#### Integration of mesoscale cortico–cortical delays

Communication between cortical areas occurs with finite transmission delays that differ for ascending and descending pathways. Feedforward signals from Layer 4 reach the soma of Layer 2/3 pyramidal neurons via short, fast axonal routes, whereas feedback signals traverse longer, polysynaptic pathways to terminate in Layer 1 on apical dendrites. As a result, feedback arrives later in physical time than feedforward input. During active perception, however, the cortex must compare both streams synchronously. One proposed solution is that higher cortical areas generate time-advanced predictions that anticipate the sensory state by *τ* milliseconds [41]. In this scheme, the delayed feedback carrying these predictions arrives just in time to coincide with the corresponding sensory evidence at the soma.

In the model described here, the feedback current reaching apical dendrites, *I*_fb_(*t*), represents the forecasted feedforward input *I*_ff_ (*t* + *τ*), such that both signals are temporally aligned when compared within the same neuron [42]. The soma, therefore, receives the actual sensory evidence, while the apical dendrite carries a prediction of that evidence at the current moment. A match preserves somato-dendritic balance, whereas any deviation, arising from an inaccurate or mistimed forecast, produces a transient imbalance that is expressed as a prediction-error spike. This computation remains gated by PV- and SOM-mediated inhibition and can be transiently released by VIP-mediated disinhibition when prediction confidence is low or mismatch uncertainty is high. In dynamic sensory contexts, this delay-compensated mechanism predicts that apical dendritic activity should lead somatic spiking by approximately *τ*, reflecting anticipatory coding. Perturbations that alter conduction velocity, interneuron timing, or temporal precision of feedback extrapolation should increase VIP-mediated disinhibition and enhance superficial-layer mismatch responses. Measuring laminar phase relationships between feedback and feedforward activity under controlled timing manipulations could provide a direct empirical test of this hypothesis.

#### Biologically motivated extensions

Future work should incorporate synaptic plasticity mechanisms to enable the adaptive refinement of prediction-error computation. Spike-timing-dependent plasticity (STDP), inhibitory scaling, or homeostatic regulation could allow the circuit to tune its connectivity based on sensory experience, increasing precision and robustness to noise [43, 44]. Another promising extension involves incorporating behavioral and motor-related modulation. Prediction-error responses in Layer 2/3 are known to depend on visuomotor context and internal state [5, 12]. Including context-dependent feedback or locomotion-related gain modulation would help explain how mismatch signals are flexibly gated in active animals.

#### Model-based predictions for experimental validation

The model generates several experimentally testable predictions regarding the cellular mechanisms of bidirectional prediction-error computation and inhibitory control:

1. **Bidirectional prediction-error coding under matched input.** When sensory input matches internal prediction, both PE^+^ and PE^−^ populations should remain near silent, maintaining a near-zero E–I balance index *β*across somatic and dendritic compartments. Simultaneous intracellular measurements of excitatory and inhibitory currents under perfectly predicted conditions could validate this balanced quiescence.
2. **Disinhibitory gating of negative prediction errors.** Activation of VIP interneurons is required to enable PE^−^ responses when feedback predictions are present in the absence of feedforward input. Inactivation of VIP should abolish these responses by reinstating dendritic inhibition, whereas activation should restore them. Optogenetic perturbations of VIP neurons during feedback-only mismatch would directly test this gating mechanism.
3. **Feature-selective prediction errors inherited from feedforward tuning.** The model predicts that prediction-error neurons preserve orientation and phase selectivity inherited from feedforward encoding in Layer 4. Even without synaptic learning, this fixed-weight architecture produces feature-specific mismatch responses. Measurements of feature-selective error activity in superficial layers would test this emergent tuning feature of the model.
4. **Dynamic inhibitory coordination maintains stability.** Reciprocal interactions between SOM and PV interneurons are predicted to maintain stable network activity during error signaling. Two-photon calcium imaging or targeted optogenetic perturbations of these inhibitory subtypes during mismatch paradigms could test the predicted inhibitory coordination that prevents runaway excitation or suppression in this model.

Together, these experiments would provide experimental evidence that tests features of this model framework proposed in this study, including that superficial cortical circuits compute bidirectional prediction errors through dynamically gated disinhibition, as well as feature-selective feedforward tuning, and balanced inhibitory coordination. This framework, therefore, forms a mechanistic bridge between bottom-up sensory encoding in Layer 4 and top-down prediction generation in deeper layers, offering a biologically grounded pathway toward a complete hierarchical model of cortical inference.

## 5 Acknowledgments

ANB and HM acknowledge support by the Australian Government through the Australian Research Council’s Discovery Projects funding scheme [DP220101166].

EN acknowledges support from a Melbourne Research Scholarship, and the Diane Lemaire and Dee & John Collier Travel Scholarships at the University of Melbourne.

## Appendix. Supplementary Methods

### A. Input preprocessing and LGN pathways

Visual input was generated from small grayscale image patches. Images were first whitened in the Fourier domain following Olshausen & Field (1996) [45] and Atick & Redlich (1992) [46], using the radial response

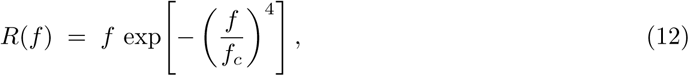

with *f_c_* = 200 defining the characteristic spatial frequency. After whitening, contrast was normalized, and each patch was convolved with an isotropic Gaussian kernel to approximate spatial integration in early vision (center–surround/LGN-like smoothing; Rao & Ballard (1999) [1]).

The smoothed signal was decomposed into ON and OFF channels by local contrast polarity: pixels above the local mean contributed to ON, and pixels below the mean to OFF, reflecting the sign-segregation of luminance pathways in the retina and lateral geniculate nucleus (LGN) Schiller (1982) [47]; Reid & Alonso (1995) [48]. Each channel was mapped onto a grid of input units ( 16 × 16) and converted into independent Poisson spike trains with mean rates *λ*_ON_ and *λ*_OFF_, plus tonic background activity of 17 Hz (ON) and 8 Hz (OFF). The resulting spike trains, *S*_LGN_(*t*), served as the feedforward drive to cortical Layer 4.

### B. Gabor filters and feature-aligned connectivity

Each excitatory neuron was assigned a Gabor receptive field:

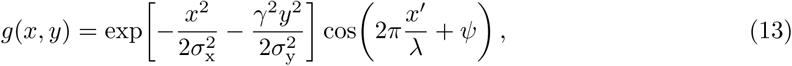

with rotated coordinates

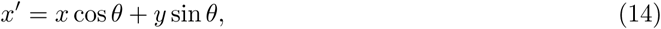

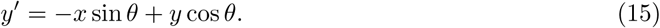

Parameters (*θ, ψ, λ, γ*) were sampled from biologically observed distributions in V1 Jones & Palmer (1987 [49]; Ringach et al. (2002) [50]. The parameters used in this study are summarised in Table 4.

**Table 4.** Gabor filter parameters for a 16 × 16 pixel receptive field with 16 pixels per degree.

| Parameter | Description | Symbol | Range / Value |
| --- | --- | --- | --- |
| Orientation | Preferred orientation of receptive field (12 evenly spaced values) | $\theta$ | $0^\circ$–$180^\circ$ in $15^\circ$ steps |
| Phase | Spatial phase offset for ON/OFF alignment | $\psi$ | $\{0^\circ, 90^\circ, 180^\circ, 270^\circ\}$ |
| Spatial frequency | Wavelength tuned to midrange visual frequency | $\lambda$ | 0.8 cycles/deg ($\approx 19.6$ px) |
| Envelope width (x/y) | Gaussian width of receptive-field envelope | $\sigma_x, \sigma_y$ | $0.17^\circ$ ($\approx 2.67$ px) |
| Aspect ratio | Reflects elongation variability of simple-cell receptive fields | $\gamma$ | 1.7–5 |

#### Parameter initialization

The receptive-field parameters were fixed across simulations to maintain controlled feature coverage. Twelve orientations (0°–180°) were uniformly distributed to emulate the diversity of orientation columns in V1. Each orientation group contained an equal number of neurons, with random phase offsets selected from the discrete set {0°, 90°, 180°, 270°} to represent contrast-polarity variants. Spatial frequency and envelope widths were chosen based on empirical tuning peaks in primate V1, and the aspect ratio *γ* was drawn from a uniform distribution to reproduce observed variability in receptive-field elongation. This initialization yields a heterogeneous but balanced population covering the principal dimensions of orientation, phase, and spatial frequency selectivity.

#### Link to predictive-coding formulation

In the original formulation of Boerlin et al. (2013) [14], each neuron contributes to the population code through an abstract decoding weight vector that determines how its spikes reconstruct the stimulus. Here, we instantiate these decoding weights using biologically grounded Gabor filters, such that each neuron’s receptive field directly defines its decoding vector. This implementation extends the predictive-coding framework by embedding realistic V1 receptive-field structure within the decoding basis, following the approach developed for Layer 4 of V1 in Nemati et al. (2025) [51]. Accordingly, the synaptic weight between neurons *i* and *j* is computed as the outer product of their receptive fields:

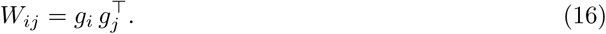

This construction ensures stronger coupling between neurons with similar orientation and phase preferences, producing feature-aligned excitation and inhibition consistent with columnar organisation in primary visual cortex. The predictive-coding dynamics operating on these structured weights are described in the following subsection.

### C. Balanced spiking predictive-coding formulation (Boerlin et al. (2013))

#### Decoding weights and outer-product connectivity

The network represents a time-varying sensory variable *x*(*t*) and its internal estimate *x*^(*t*). Following Boerlin et al. (2013) [14], the estimate is obtained by a leaky linear readout of the population:

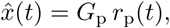

where *G*_p_ is the matrix of *decoding weights* (each column is a neuron’s decoder), and *r*_p_(*t*) are exponentially filtered spike trains of the neurons. Each neuron spike therefore adds its neuron’s decoding vector to *x*^(*t*), contributing a small correction toward the target signal *x*(*t*).

The network minimizes the instantaneous reconstruction error ||*x*(*t*) − *x̂*(*t*)||^2^ by spiking whenever this error exceeds a neuron’s threshold. To ensure efficient coding, recurrent couplings must cancel redundant contributions between similarly tuned neurons. This cancellation is achieved when the fast synaptic couplings are proportional to the pairwise overlaps of decoding vectors:

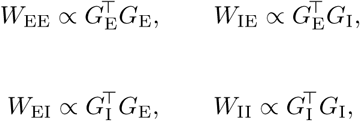

where *W*_EE_, *W*_IE_, *W*_EI_, and *W*_II_ denote fast recurrent synaptic weight matrices between excitatory (E) and inhibitory (I) populations. Feedforward connections are aligned with the same decoding basis, *W*_FF_ ∝ *G*_E_, ensuring that sensory evidence arrives in the same representational space as the population readout.

Intuitively, if two neurons have highly overlapping decoding vectors, a spike from one will transiently suppress the other’s prediction error via the recurrent term. This outer-product connectivity thus enforces local balance and decorrelation in feature space.

#### Dynamics

Membrane potentials represent local prediction errors, integrating feedforward drive and recurrent feedback.

For excitatory neurons:

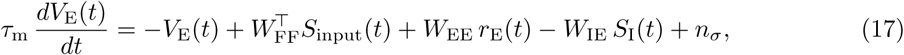

and for inhibitory neurons:

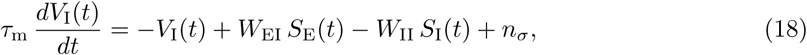

where *V*_E_(*t*) and *V*_I_(*t*) are membrane potentials, *S*_E_(*t*) and *S*_I_(*t*) are spike trains of excitatory and inhibitory neurons, *S*_input_(*t*) are feedforward input spikes, *τ*_m_ is the membrane time constant, and *n_σ_*is zero-mean Gaussian noise.

A neuron emits a spike when *V_i_*(*t*) *>* Θ*_i_*, after which *V_i_* is reset to *V_r_*for a refractory period. Synaptic currents are exponentially filtered with time constants *τ*_E_ (excitatory) and *τ*_I_ (inhibitory), maintaining fast excitation–inhibition balance. Under these dynamics, voltages continuously track projection-specific prediction errors, and each spike implements a discrete correction that reduces ||*x*(*t*) − *x̂*(*t*)||^2^.

### D. Layer 4 network providing feedforward input to Layer 2/3

Layer 4 (L4) supplies the present model with a sparse, feature-structured spike code and tightly balanced excitatory–inhibitory (E–I) drive. We use the same L4 as in our companion work Nemati et al. (2025) [51]: a recurrent spiking network with a canonical 4:1 E:I ratio obeying Dale’s law. Feedforward input consists of ON/OFF LGN-like Poisson spike trains generated from whitened, contrast-normalized image patches (Appendix A); excitatory neurons carry Gabor-parameterized decoding weights (Appendix B), so that ON/OFF afferents project in a feature-aligned basis. Recurrent couplings are proportional to decoder overlaps (outer-product structure), implementing the spike-based predictive-coding dynamics of Boerlin et al. (2013) [14] with biophysically plausible synaptic filters and refractory periods (Appendix C).

In this regime, Layer 4 exhibits hallmark features of balanced cortical dynamics. (i) **Excitatory–inhibitory balance:** Individual excitatory and inhibitory neurons maintain near-proportional excitation and inhibition, with slopes close to 1, strong negative correlations between excitatory and inhibitory input currents, and a mean balance index B ≈ 0.01 (ratio of residual net current to total input magnitude; B ≪ 0.1 indicates tight balance). (ii) **Asynchronous–irregular regime:** Spike trains are irregular and weakly correlated across neurons, with an inter-spike-interval coefficient of variation (ISI CV) of approximately 1.05–1.10 and near-zero pairwise synchrony, consistent with in-vivo recordings of Layer 4 in primary visual cortex. (iii) **Emergent feature selectivity:** Without any learning or plasticity, excitatory neurons develop orientation and phase tuning: for drifting gratings, 43.6% of excitatory cells show a strong orientation bias index (*OBI >* 0.7) and 81.7% show a strong phase bias index (*PBI >* 0.9). Inhibitory neurons display broader tuning, reflecting their integrative role. Both populations exhibit simple-cell–like modulation ratios (*F*_1_*/F*_0_ *>* 1). (iv) **Contrast gain with preserved balance:** When stimulus contrast increases, excitatory and inhibitory population firing rates rise proportionally, and their scaled temporal profiles remain tightly matched, with a mean-squared-error (MSE) between them below 0.005. (v) **Robust decoding:** Linear reconstruction of the input from excitatory population firing rates remains accurate even when up to roughly 50% of image pixels are replaced with random noise, demonstrating resilient sensory encoding under degraded input.

For the present study, Layer 4 provides spike trains from its excitatory population as the feeforward input to Layer 2/3. No top-down feedback targets Layer 4 in this model, consistent with anatomical evidence that corticocortical feedback and higher-order predictions primarily terminate in supragranular (Layer 2/3) and infragranular (Layer 5/6) compartments rather than the granular layer. Accordingly, Layer 4 operates here as a purely feedforward stage that transforms LGN drive into feature-selective spike patterns, supplying sensory evidence to the prediction-error circuit in Layer 2/3. Comprehensive implementation details are available in Nemati et al. (2025) [51].

Together, Appendices A–D provide a complete specification of the feedforward pathway used in the present Layer 2/3 model, from image preprocessing to feature-selective spike generation in Layer 4.

